# High-throughput single molecule microscopy with adaptable spatial resolution using exchangeable oligonucleotide labels

**DOI:** 10.1101/2024.10.30.621026

**Authors:** Klarinda de Zwaan, Ran Huo, Myron N.F. Hensgens, Hannah Lena Wienecke, Miyase Tekpinar, Hylkje Geertsema, Kristin Grussmayer

## Abstract

Super-resolution microscopy facilitates the visualization of cellular structures at resolutions approaching the molecular level. Especially, super-resolution techniques based on the localization of single-molecules have relatively modest instrument requirements and are thus good candidates for adoption in bioimaging. However, their low throughput nature hampers their applicability in biomolecular research and screening. Here, we propose a workflow for more efficient data collection, starting with scanning of large areas using fast fluctuation based imaging, followed by single-molecule localization microscopy of selected cells. To achieve this workflow, we exploit the versatility of DNA oligo hybridization kinetics with DNA-PAINT probes to tailor the fluorescent blinking towards high-throughput and high-resolution imaging. Additionally, we employ super-resolution optical fluctuation imaging (SOFI) to analyze statistical fluctuations in the DNA-PAINT binding kinetics, thereby tolerating much denser blinking and facilitating accelerated imaging speeds. Thus, we demonstrate 30-300-fold faster imaging of different cellular structures compared to conventional DNA-PAINT imaging, albeit at a lower resolution. Notably, by tuning image medium and data processing though, we can flexibly switch between high-throughput SOFI (scanning a FOV of 0.65*mm* × 0.52*mm* within 4 minutes of total acquisition time) and super-resolution DNA-PAINT microscopy and thereby demonstrate that combining DNA-PAINT and SOFI enables to adapt image resolution and acquisition time based to the imaging needs. We envision this approach to be especially powerful when combined with multiplexing and 3D imaging.

## Introduction

Fluorescence microscopy has elucidated important new insights in cellular processes over the past decades. The recent establishment of super-resolution microscopy methods have further pushed the boundaries of fluorescence microscopy to facilitate the visualization of cellular structures, such as nuclear pores and the neuronal cytoskeleton, at resolutions close to the molecular level^1^. A particular promising SMLM method is DNA-PAINT (Point Accumulation for Imaging in Nanoscale Topography) since it can achieve the highest localization accuracy while still posing only modest hardware requirements. DNA-PAINT uses transient hybridization events between fluorophore-coupled single-stranded DNA oligonucleotides (imagers) and target-associated complementary DNA oligonucleotides (docking strands) that can be bound repetitively^2^. The temporary immobilization of imagers yields a distinct fluorescent blink that can be computationally localized with an accuracy of a couple of nanometers. The oligo blinking kinetics are highly programmable by modulating DNA hybridization parameters through sequence design and buffer composition to obtain only a subset of the target-bound oligos to be hybridized to fluorescent imagers, thereby separating single blinking events in space and time. Localizations of tens of thousands individual fluorescent blinks then render a super-resolved image. Compared to other SMLM techniques including dSTORM or PALM where photoswitching between on- and off-states are tuned by laser excitation, or chemical reagents, DNA-PAINT blinking events are solely determined by DNA oligo hybridization kinetics and thus uncoupled from fluorophore photophysics allowing for the use of the brightest, photostable organic dyes. DNA-PAINT also facilitates multiplexed imaging of different cellular structures in a single wavelength through use of different imager-binder pairs sequentially, thereby avoiding chromatic aberration.

Despite the many advantages of DNA-PAINT, this method is still sparsely deployed, arguably due to its low data throughput. The fluorescent signal of immobilized imagers needs to be segregated from diffusive background fluorescence by utilizing long exposure times of several hundreds of ms, making DNA-PAINT the slowest super-resolution detection method. In addition, it is not straightforward to know prior to data acquisition and analysis if the observed cells are tangible, leading to notable time spent on measuring intangible data sets. This is a significant problem for expanding this promising visualization technique to advance our understanding of the nanoscopic organization of cellular molecules and is exacerbated in 3D imaging.

Here, we aim to improve the detection speed by combining DNA-PAINT with fluctuation-based imaging using super-resolution optical fluctuation imaging (SOFI) and facilitate high throughput super-resolution imaging. While both rely on stochastic blinking of single fluorophores, SMLM and SOFI differ in the mechanisms exploiting information below the diffraction limit. SOFI uses higher-order statistical analysis of time series of blinking molecules (i.e., often a fluctuating signal from several overlapping fluorophores) to reconstruct super-resolution images, and avoids the need to localize individual fluorophores. Correlations in the blinking signal allow us to perform spatiotemporal cumulant analysis with a moderate amount of frames to construct images with a super-resolution point-spread function raised to the power of the cumulant order n. SOFI is relatively insensitive to background signal, allowing for higher labeling densities, higher blinking on-time ratios, lower signal-to-noise and reduced acquisition times than applicable for DNA-PAINT. Previous work by Glogger et al.^3^ showed that SOFI can be combined with exchangeable labels using standard DNA-PAINT to eliminate photobleaching effectively. Building on this, we utilize the programmability of DNA-PAINT kinetics and combine it with advanced SOFI processing to significantly speed up super-resolution imaging.

By tuning DNA-PAINT kinetics complemented with SOFI data processing, we establish super-resolution whole-cell imaging of microtubules and mitochondria with 2nd order SOFI in 5 seconds (500 frames), and up to 4th order SOFI in 50 seconds (5000 frames), which is 30-300 times faster compared to SMLM data acquisition. Additionally, our approach allowed us to successfully achieve high-order SOFI up to 6th order, with a resolution of 75 nm. Moreover, we also demonstrate that we can effectively switch between two super-resolution modalities - SOFI and SMLM. As a consequence, high-throughput imaging by SOFI can provide a quick sample overview at improved resolution and deliver the necessary optical sectioning for e.g. identification of rare phenotypes. Subsequent SMLM imaging of selected cells allows the ultimate zoom-in at the highest resolution by leveraging the full resolving power of DNA-PAINT labels.

## Results

### High-order SOFI using DNA-PAINT

In this study, we explore the synergistic integration of DNA-PAINT and SOFI to accelerate super resolution imaging. SOFI capitalizes on the fluctuations of fluorescent signals to achieve resolution beyond the diffraction limit (Supplementary Note 1)^4^. A better measurement of blinking statistics will thus lead to better SOFI signal, which is critical to exploit higher order SOFI and thus higher spatial resolution. We focus on increasing the frequency of fluorescent fluctuations for SOFI analysis by adapting the binding kinetics of DNA-PAINT probes (Fig 1).

**Figure 1.**
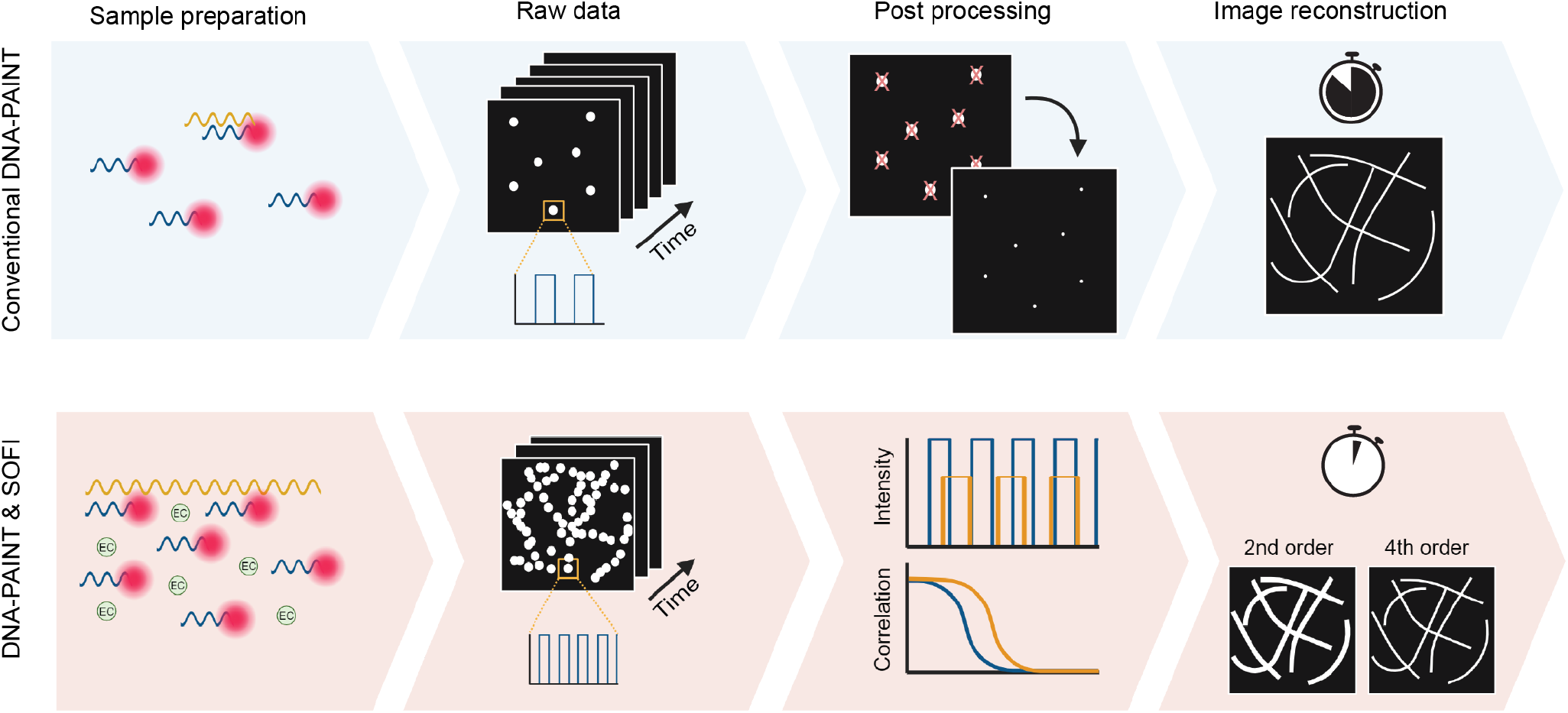
Overview of integration between DNA-PAINT and SOFI. The upper panel illustrates the conventional DNA-PAINT workflow, where a time series is recorded, individual emitters are localized with high accuracy, and a super resolved image is reconstructed. The lower panel demonstrates the integration between DNA-PAINT and SOFI. Here, optimized sample preparation is achieved through sequence design and buffer conditions to increase the frequency of fluorescent fluctuations, giving a higher emitter density per frame. SOFI calculations are then performed using a pixel-by-pixel cross-cumulant algorithm combined with brightness linearization, producing high-order SOFI images.

In our experiments, we used speed optimized DNA sequences with periodic sequence motifs (5xR1, see table 1) that have shown to provide a 5-fold increase in binding frequency^5^ compared to standard DNA-PAINT probes. In addition, we work with high imager strand concentrations in the nanomolar range to further raise the probability of hybridization (see Supplementary Note 2). Photobleaching does not play a role in our experiments (see Supplementary Figure 8) since imager strands are exchangeable and can be continuously replenished from a practically infinite buffer reservoir. Taken together, those experimental refinements allowed us to achieve up to 6th order SOFI of microtubules in fixed COS-7 cells shown in Figure 2. SOFI calculations were performed using a cross-cumulant-based algorithm with postprocessing including deconvolution and brightness linearization which is essential for high cumulant orders^6,7^ (see Supplementary Note 1), thereby overcoming the limitations of Glogger et al. that obtained up to third-order reconstructions using conventional DNA-PAINT imagers.

**Table 1.**
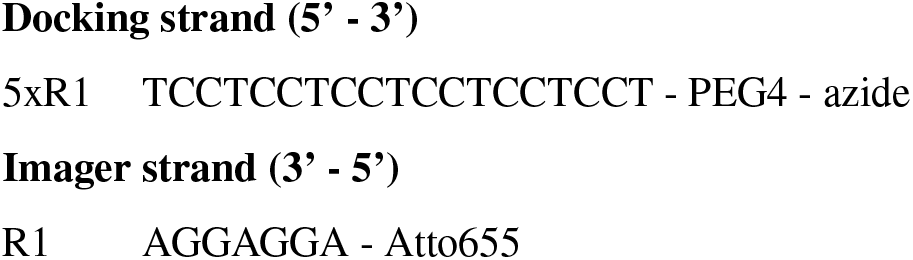
Docking site sequences and modifications for nanobody conjugation and corresponding imager strand sequences.

**Figure 2.**
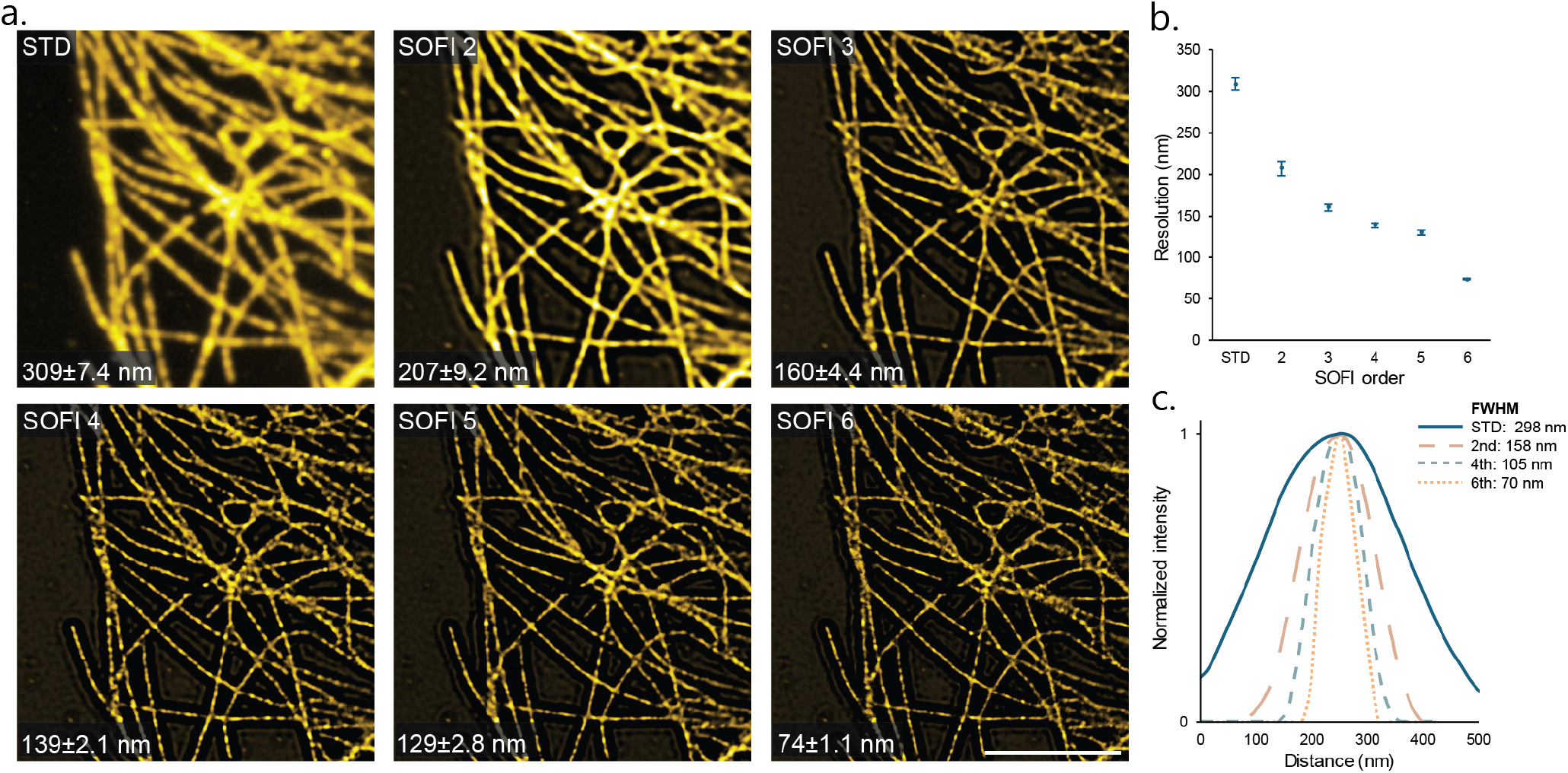
High-order DNA-PAINT SOFI reconstructions. COS7 cells stained for microtubules with repeating docking sequence with resolution increasing for increasing cumulant order. (a) Close-ups of diffraction limited standard deviation of the image sequence, 2nd, 3rd, 4th, 5th and 6th order SOFI (scale bar 5 μm). (b) Resolution estimate by decorrelation analysis for three different cells (average ± standard deviation)^8^. (c) Microtubule cross-section intensity profile for the standard deviation, 2nd, 4th and 6th order SOFI and corresponding FWHM measurements.

SOFI effectively suppresses background noise and improves optical sectioning; this is already apparent in second-order reconstructions in Figure 2. In contrast, the widefield standard deviation image shows prominent out-of-focus background, preventing the clear distinction between adjacent microtubule filaments. Each successive order n in SOFI contributes to resolving finer structural details, providing theoretically an up to *n*-fold resolution enhancement with subsequent deconvolution^9^. To quantify the SOFI results and confirm the expected resolution increase with successive orders, the spatial resolution is estimated using image decorrelation analysis^8^ in Fig 2b. The resolution enhancement is in good agreement with theoretical predicitions. Specifically, for 6th order SOFI we achieve a remarkable resolution of approximately 75 nm. As a second metric for resolution, the intensity profile across the microtubule axis is quantified (Fig 2c). These results are consistent with decorrelation analysis, showing an increase in resolution with higher orders. For 6th order, the mean diameter of the microtubule (FWHM) is 70 nm. In addition, we show in Supplementary Fig **??** mitochondrial structures that are also resolved up to 6th order SOFI with the expected resolution enhancement, demonstrating the versatility of our approach.

Higher-order statistical analysis is challenged by the photophysical properties of the fluorphores used, limiting the usage of most fluorophores. First, the ideal fluorophore for SOFI should be photostable^4,10^. Photobleaching, a correlated phenomenon, will affect the results and would need to be corrected for in the analysis. DNA-PAINT excels in this regard, as its blinking events are decoupled from the inherent fluorophore photophysics and it is considered resistant to photobleaching^3^. Moreover, calculating higher-order cumulants requires well sampled statistics and homogeneous fluorescence blinking behavior^6^; both is the case for DNA-PAINT labels with fast fluctuations and uniform, programmed oligonucleotide binding-unbinding kinetics. Many fluorophores, however, exhibit inhomogeneous blinking during the measurement time, which limits their utility for analysis beyond 2nd or 3rd order SOFI and can lead to artifacts. In addition, DNA-PAINT probes enable the use of the brightest organic dyes. Altogether, DNA-PAINT labels tuned towards high fluctuations are particularly well-suited for high-order SOFI reconstructions due to their exceptional blinking behaviour.

### Reducing the acquisition time

Next, our objective was to enhance the fluctuations to a level that allows us to increase the sampling rate and reduce the number of frames required, all while maintaining high SOFI quality. To achieve this, we optimized our imaging buffer (Supplementary Note 2) by using even higher imager strand concentrations (with the periodic sequence motif docking strand) to decrease the off-time. At the same time, we add the small molecule ethylene carbonate (EC) to reduce the on-time by destabilizing the DNA duplex, leading to more pronounced intensity fluctuations. These optimizations enabled us to measure at a higher frame rate of 100 Hz due to the greater frequency of binding and unbinding events.

As a result, we reduced the minimal acquisition time for 2nd order SOFI to only 5 seconds (or 500 frames) and for 4th order to 50 seconds (or 5000 frames) (Fig 3a,b), which falls within previously reported ranges for other fluorophores^6^ and is 25-fold faster than the DNA-PAINT SOFI acquisitions of Glogger et al. These achievements are validated through a qualitative assessment of structural continuity and absence of artifacts for different acquisition times while the resolution is preserved (Supplementary Fig **??**). Additionally, a pixel-wise SNR estimation based on a statistical approach known as jackknife resampling was performed^11^ to quantify the SNR of the SOFI images for different acquisition times (Fig 3d).

**Figure 3.**
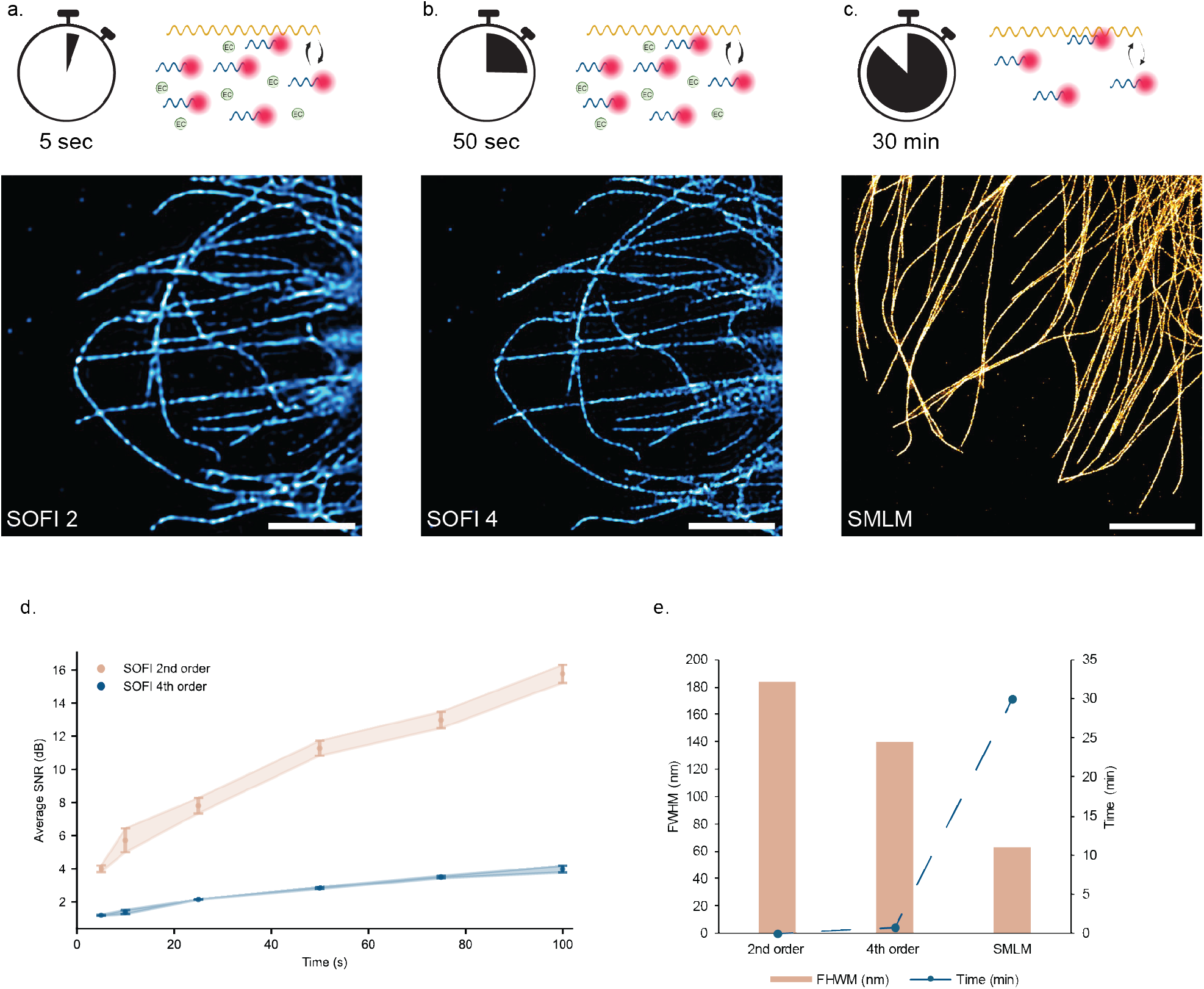
Minimum acquisition time for SMLM and SOFI DNA-PAINT. (a, b) SOFI DNA-PAINT reconstructions of fixed microtubules in COS7 cells labeled with repeating docking sequence obtained with a minimal number of frames and with a frame rate of 100 Hz (scale bar 5 *μ*m, Supplementary Fig **??**a,b for qualitative assessment of minimal frame acquisition). (c) SMLM DNA-PAINT reconstructions obtained with a minimal number of frames and with a frame rate of 10 Hz (scale bar 5 *μ*m, Supplementary Fig **??**c for qualitative assessment of minimal frame acquisition). (d) Jacknife SNR metric for 2nd and 4th order SOFI as a function of acquisition time. (e) Average of 5 FWHM measurements of microtubule crosssections for each reconstruction correlated with the minimum acquisition time.

SMLM DNA-PAINT requires data acquisition at 10 Hz to ensure a sufficient signal-to-background ratio that enables accurate localization. Localization of single emitters was executed and we observed continuity in microtubule filaments after a time series of 30 minutes with a localization precision of 7 nm (Fig 3c). In comparison, our DNA-PAINT approach combined with SOFI analysis facilitated a 30-fold reduction in acquisition time for 4th order SOFI (50 seconds) and a 300-fold reduction for 2nd order SOFI (5 seconds). This highlights that, while SMLM achieves higher spatial super-resolution, it does so at the cost of increasing the acquisition time. Conversely, SOFI improves the acquisition speed (i.e. temporal resolution for dynamic samples) but compromises spatial resolution. This is exemplified by the intensity profile across the microtubule axis for each image (Fig 3e) which is in line with decorrelation analysis. Nevertheless, the SOFI optimized buffer allows for rapidly achieved super-resolution marking a significant advancement in imaging speed and highlighting its potential for high-content super-resolution imaging applications.

### High-throughput super-resolution imaging for screening applications

To address the inherent trade-off between spatial and temporal resolution in traditional SMLM DNA-PAINT, we developed a high-throughput screening workflow that integrates SOFI followed by SMLM. This approach leverages the fast data acquisition of SOFI to rapidly acquire super-resolution images, which are then used to guide subsequent imaging for SMLM. The sample remains unchanged, with only the buffer conditions modified to achieve the sparse blinking necessary for single-molecule localization (Fig 4a). The drastic decrease in image acquisition time enables high-throughput super-resolution imaging by SOFI directly. To demonstrate, we acquire a 0.65*mm*× 0.52*mm* area by subsequently scanning partially overlapping FOVs in a 4 × 9 grid, screening approximately >40 cells (Fig 4a). At each grid position we cover a FOV of 84μ*m* by 150μ*m* and image for 500 frames at 10 ms exposure time using homogeneous illumination. The total imaging time is thus 3 minutes plus additional 1 minute of stage movement and data saving time using an automated multi-position imaging protocol. Second-order SOFI reconstructs the microtubule network for the whole stitched FOV at a two-fold resolution enhancement (Fig 4a).

**Figure 4.**
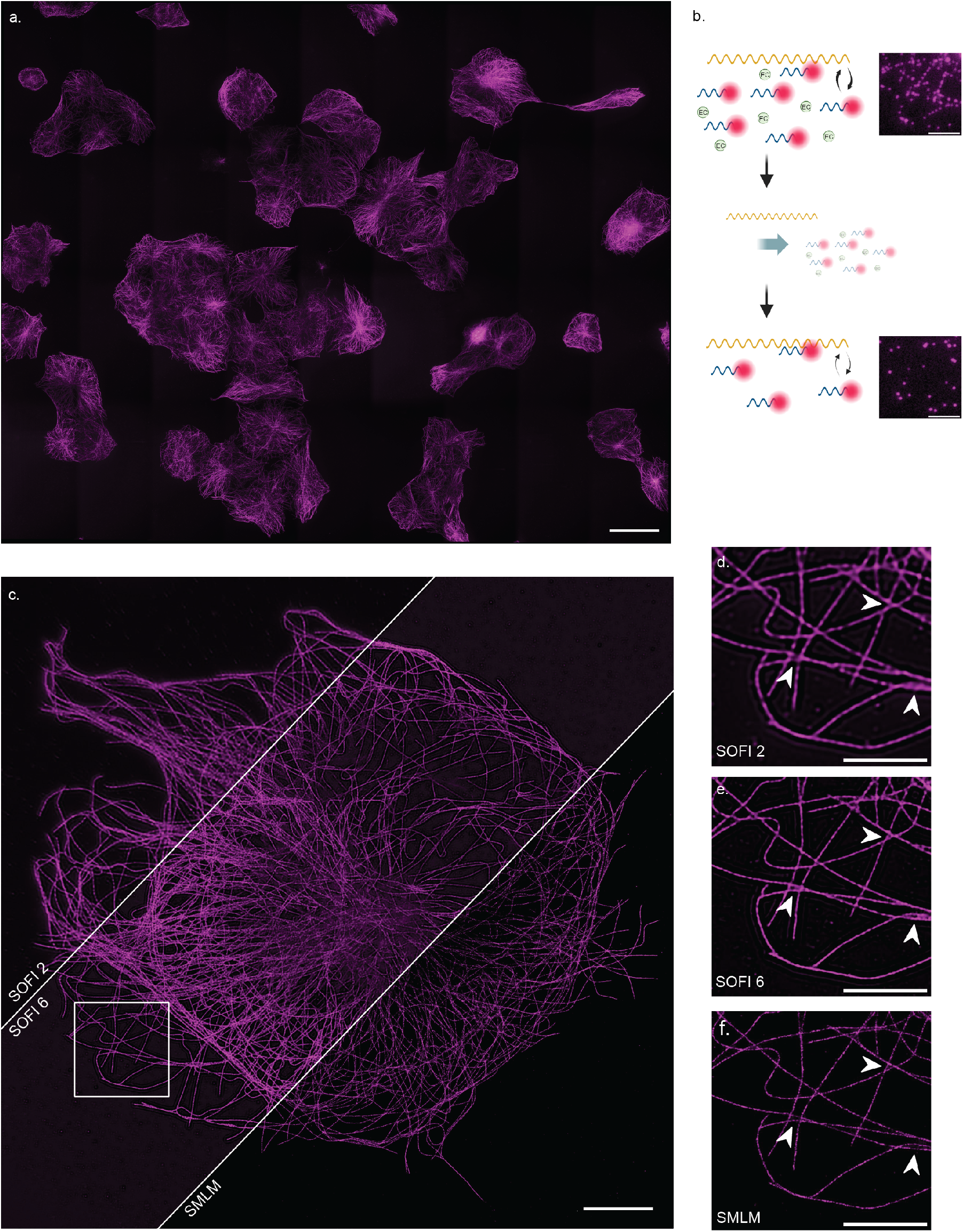
Proof of principle: High-throughput SOFI followed by SMLM. (a) High-throughput 2nd order SOFI imaging of a 0.65*mm*× 0.52*mm* FOV containing about 20 cells (scale bar 50 *μ*m). (b) Scheme showing the workflow: start imaging with a buffer optimized for fast SOFI, wash away buffer components, introduce low concentration of imager strands. Close up of raw data frames used for SOFI (top) and SMLM (bottom) reconstructions of COS7 cells stained for microtubules (scale bar 5 *μ*m). (c) SOFI and SMLM reconstructions of the same FOV (scale bar 10 μm). (d, e, f) Close-ups of corresponding SOFI and SMLM reconstructions as indicated. Arrows indicate areas where improvement of resolution is visible (scale bar 5 *μ*m).

Subsequently, we image one field of view (about 100 × 100 μm) first in the SOFI regime, followed by the SMLM regime (Fig 4c). To reconstruct a 2nd order SOFI image (Fig 4d), we need 75 seconds of acquisition time. The longer acquisition time compared to the results shown in Figure 3 was due to our use of a reduced imager strand concentration. This adjustment was necessary to ensure thorough washing away of all imager strands within a reasonable timeframe. To obtain sufficient blinking statistics also for higher-order SOFI, we extend imaging to a total of 4 minutes. This allows us to achieve high-quality 6th order SOFI, resulting in the expected resolution improvement (Fig 4e). After washing, we acquired SMLM data for a minimum of 25 minutes to reconstruct continuous microtubules with a localization precision of 8.35 nm (Fig 4f).

This workflow optimizes the imaging process, making it suitable for applications that require very high localization precision and spatial resolution, but also benefit from fast screening and efficient data acquisition. SOFI inherently provides optical sectioning and background reduction thereby facilitating further downstream image processing. For instance, it will be particularly advantageous for screening a large number of samples or cells to identifying those suitable for downstream analysis or even to recognize biologically rare events. Moreover, this approach is fully compatible with traditional DNA-PAINT multiplexing approaches and 3D imaging, offering the possibility to multiply the information content. Overall, the integration of SOFI and SMLM in our single molecule-based super-resolution workflow significantly improves imaging efficiency without compromising resolution quality.

## Discussion

In summary, our results showed the compatibility of exchangeable labels with two super-resolution techniques, DNA-PAINT and SOFI, and the advantage of significant imaging speed increase when combining them. We also demonstrated spatial resolution tuning in high throughput imaging with our method.

While both rely on the stochastic blinking of single fluorophores, DNA-PAINT and SOFI differ in the mechanism for extracting information below the diffraction limit. SOFI gains resolution enhancement from statistical analysis of detectable fluorescence fluctuations. The quality of SOFI images depends on the effective contrast between on- and off-states, the SNR of acquired images, and the sampling of the blinking kinetics. Homogeneous blinking kinetics are beneficial for SOFI and low photobleaching ensures that the spatial and temporal correlations analyzed in SOFI arise from stochastic fluorescence fluctuations^6^. Meanwhile, DNA-PAINT is a single-molecule localization technique that decouples blinking events from fluorophore photophysics with exchangeable and reversibly binding labels^2,12^, providing flexibility to modulate the blinking kinetics through various pathways. In addition, photobleaching is largely circumvented due to the abundant imager strands that are replenishing from the buffer. Therefore we see the potential in DNA-PAINT probes as a great supplement for high spatiotemporal resolution imaging using SOFI.

In this work, we used the exchangeable oligonucleotide-based probes and first sped up the blinking kinetics for SOFI using repeating sequences on the docking strand, which has been shown to increase the binding events frequency^5^. The highly-correlative fluorescence fluctuations resulting from frequent binding and unbinding events at high imager concentrations is crucial for high-order SOFI analysis. We achieved the first successful 6th order SOFI reconstruction of cellular structures with DNA-PAINT probes. This required the use of post-processing algorithms including deconvolution and brightness linearization, resulting in the improvement of the 6th order SOFI resolution up to 70 nm. Compared to previous work utilizing exchangeable nucleotide based probes for SOFI^3^, this has increased the resolution enhancement by approximately three-fold, and is in line with the best resolution reported in imaging continuous cellular structures for SOFI, which is 50-60 nm at 6th order^6^.

Similar to other SMLM techniques, conventional DNA-PAINT suffers from a long acquisition time. Advances in speeding up DNA-PAINT have been focusing on accelerating the blinking kinetics, i.e. shortening both the on- and off-time of blinking fluorophores. However, shorter on- and off-time translate to more fluorophores present in each frame, which poses challenges for SMLM due to the overlapping PSFs in a dense frame that lead to image artifacts. In addition, the localization precision suffers from shorter acquisition times due to a reduced signal-to-background ratio. SOFI eliminates the requirement of sparse distinguishable fluorophores, thereby opening up more blinking kinetics space for imaging speed up. We combined several strategies to increase the blinking frequency. Next to using repeating sequences on the docking strand, we simply increased the imager strand concentration to increase the binding events frequency and to decrease the off-time. We also added ethylene carbonate to the imaging buffer^12^, which destabilizes DNA duplex in order to increase the dissociation rate and to decrease the on-time. This resulted in a 2nd order SOFI image of the microtubule network in cells within only 5 seconds, or 500 frames at 100 Hz. Our data acquisition is 25-fold faster compared to Glogger et al., where the total acquisition time amounts to 125s using the standard P1 and P4 sequences. This improvement is in the same order of magnitude as a novel SOFI pre-processing algorithm that modifies the SNR and SNB^13^ for dense fluorescent protein data through deconvolution. Obtaining more frames for as long as 50 seconds lead to 4th order SOFI reconstructions in our measurements, 30-fold faster than a typical SMLM acquisition.

Compared to other methods to accelerate DNA-PAINT, for example, argo-PAINT^14^ and FLASH-PAINT^15^, we avoided adding additional protein or nucleic acids or greatly extending the imager strand length. Our approach could also be combined with other factors in the buffer affecting the nucleic acid binding kinetics, such as salt concentration and temperature^16^. The limitation for our method for more acceleration is mainly the high background signals at higher imager strand concentration that eventually compromises the SNR, even though SOFI intrinsically suppresses the non-correlative background noise. We used TIRF or HILO illumination for our images to provide extra optical sectioning that facilitated the higher order SOFI. The upper limit of acceleration supported by increased blinking kinetics depends on the structures of interest and the docking strand labeling efficiency. The recent fluorogenic and self-quenched imager strands^17,18^ can further help to reduce background and expand applications in thick samples. 3D SOFI where different z-positions are imaged at the same time, for example through multiplane splitting^9^, can further increase the throughput of the approach.

The optimization of spatial and temporal resolution of DNA-PAINT-SOFI not only increases the imaging speed at high resolution and high throughput, but can also function as a useful tool for fast high-content screening of samples at a moderate resolution enhancement. The drastic reduction in acquisition time allowed for a 4-minute imaging with 2-fold resolution enhancement, scanning through a total FOV of 0.65*mm*× 0.52*mm*. We demonstrated that we can conveniently switch from SOFI conditions to SMLM with localization precision of few nanometers, simply by modifying the buffer composition, i.e. by lowering the imager strand concentration via a microfluidic system. The resolution improvement between fluctuation-based (about 70 nm) and localization imaging (about 7 nm) in our workflow is akin to switching between confocal and STED imaging, which is routinely done to facilitate data acquisition.

Our second-order SOFI acquisition time for a single position is of similar scale as structured illumination imaging with DNA-PAINT labels^19,20^ that enables a maximum 2-fold resolution increase. Importantly, our SOFI to SMLM workflow can be carried out using a microscope with simple hardware, facilitating straightforward adoption of our proposed approach. We envision screening a large number of cells with fast SOFI, and to use e.g. machine learning algorithms to interrogate the optically sectioned and background-reduced images to identify rare phenotypes for subsequent interrogation by DNA-PAINT. Since DNA-PAINT relies on stochastically blinking single molecules, identification of full protein structures or networks, and thereby rare events, is generally hampered by time-intensive image acquisition. In fact, the continuous adjustment capability of blinking kinetics with exchangeable oligonucleotide-based probes facilitates tuning of temporal and spatial resolutions to visualize protein structures and networks from a few nanometers with SMLM to dozens with SOFI.

Finally, the method we propose is not limited to fixed cells. Novel PAINT-alike probes compatible with live cells, such as self-labeling protein tags labelled with reversible fluorescent probes^21^ offer a promising outlook for high-content live-cell super-resolution imaging. We envisioned our method contributing towards the goal of fast 3D multi-target super-resolution imaging.

## Methods

### Nanobody production

Bacterial expression plasmids pTP1122 and pTP955 and pDG02583 were a gift from Dirk Görlich (Addgene plasmid #104159; http://n2t.net/addgene:104159; RRID:Addgene_104159, Addgene plasmid #104164; http://n2t.net/addgene:104164; RRID:Addgene_104164, Addgene plasmid #104129; http://n2t.net/addgene:104129; RRID:Addgene_104129, respectively)^22^.

The anti-mouse and anti-rabbit nanobodies with protease-cleavable affinity tags and engineered cysteines, and *bd*NEDP1 protease fused to His14-MBP-*bd*SUMO were expressed in *E*.*coli* BL21(DE3)^22^. 2 liters of Luria Bertani broth (LB broth) was inoculated with 20 mL overnight culture. E.coli were grown to an OD600 between 0.4 and 0.7 before protein expression was induced by 0.5 mM isopropyl B-D-1-thiogalactopyranoside (IPTG). 4 hours after induction, cells were pelleted by centrifugation and resuspended into lysis buffer (50 mM Tris/HCL; pH 8.0, 1 M NaCl, 5mM beta-mercaptoethanol, 50 mM imidazole) and 1mM PMSF was added. Cells were lysed by sonication, and the lysate was cleared by ultracentrifugation for 30 minutes at 4^°^C (Ti45 rotor, 37,000rpm, Beckman Coulter). The proteins were purified by affinity chromatography using an ÄKTA Start (GE Healthcare ÄKTA Start). The lysate was passed through a 5 mL pre-equilibrated HisTrap HP column (Cytiva) and was washed with lysis buffer. Gradient elution was performed over 10 column volumes (CVs) with filter sterilized elution buffer (50 mM Tris/HCl; pH = 8.0, 150 mM NaCl, 5 mM beta-mercaptoethanol, 500 mM imidazole). The fractions containing nanobody were pooled and a buffer exchange to maleimide labelling buffer (MLB; 100 mM potassium phosphate buffer; pH = 7.5, 150 mM NaCl, 250 mM sucrose) using SnakeSkin dialysis tubing was performed. For the *bd*NEDP1 protease, the eluate was rebuffered to protease buffer (50 mM Tris/HCl; pH = 7.5, 300 mM NaCl, 250 mM sucrose). The protein concentration after buffer exchange was determined using a Nanodrop1000 spectrophotometer (Thermo Fisher Scientific). The nanobodies and protease were aliquoted, frozen in liquid nitrogen, and stored at -80^°^C until further use.

1 mM His-tag containing nanobodies were cleaved by 0.6 μM bdNEDP1 protease in the thermoshaker at 20^°^C and 300 RPM for 24 - 96 hours. Cleaved His-tags, His-tag containing proteases and uncleaved nanobodies were purified out of the solution by reverse affinity chromatograpy using an ÄKTA Start (GE Healthcare ÄKTA Start). The cleaved mixture was subjected to purification using a pre-equilibrated 1 mL HisTrap HP column (Cytiva). After loading the sample, the column was washed with MLB to separate and collect the unbound protein (i.e. cleaved nanobodies). Subsequently, gradient elution was conducted over 5 CVs using MLB supplemented with 500 mM imidazole. During elution, the cleaved tags and protease were collected. The purity of the nanobodies was assessed using SDS-PAGE, and the protein concentrations were measured using a Nanodrop1000 spectrophotometer (Thermo Fisher Scientific). Subsequently, the cleaved and purified nanobodies were aliquoted, frozen in liquid nitrogen and stored at -80^°^C until further use.

### Sample preparation

#### Site-specific labeling of nanobodies

A site-specific labeling protocol of the nanobodies with an azide functionalized DNA oligonucleotide was developed based on previous literature and contains a two-step reaction^12,23^. First, a DBCO-maleimide linker is conjugated to the nanobodies with engineered cysteines. Second, 5’-azide functional oligonucleotide docking strands are conjugated.

Purified and cleaved nanobodies with engineered cysteines were freshly reduced with a 30-fold molar excess of 15 mM TCEP (Carl Roth) for 30 minutes on ice. For a standard reaction, 40 μM reduced nanobody was mixed with 2mM DBCO-maleimide (Jena Bioscience) and incubated for 4 hours at 4^°^C. Unbound reaction partners were removed in two buffer exchange steps with Phosphate Buffer Saline (PBS; Gibco, ThermoFisher) using a Zeba spin desalting column (10,000 MWCO).

The protein concentration and the degree of labeling (DOL) were determined by absorbance at 280 nm for the nanobodies and 309 nm for the DBCO using a Nanodrop1000 spectrophotometer (Thermo Fisher Scientific).

Docking strand-oligonucleotides (see table 1), modified with either a 3’ or a 5’ azide moiety were synthesized by Biomers.net (Germany) and dissolved in PBS to a concentration of 5mM. For a standard reaction, 10 μM was incubated with 300 μM azide-docking strand for 30 minutes at 20^°^C at 300 rpm. Unconjugated docking strands were removed similar to the DBCO conjugation using a Zeba spin desalting column (10,000 MWCO) and protein concentration was determined by measuring the absorbance at 280 nm. The conjugated nanobodies were either stored at 4^°^C in PBS or -20^°^C in 50% glycerol.

Optimal molar ratios and reaction times for the linker conjugation were determined using maleimide-modified methoxy-polyethyleneglycol (mPEG, MW = 5000; Biopharma PEG) as a click-reaction partner. The reaction efficiency was determined using molar ratios of nanobody to maleimide-mPEG ranging from 1:10 - 1:100 and incubating for 4 to 24h. Reaction samples were quenched by denaturing and analyzed using SDS-PAGE. Similarly, optimal incubation conditions were determined for the docking strand conjugation using azide-mPEG (MW = 5000; Biopharma PEG). Molar ratios ranged from 1:1 - 1:50 and incubation times ranged from 0.5 to 24h. Additionally, the DOL was determined for the different incubation conditions.

#### Cell culture

COS-7 cells (DSMZ GmbH) were cultured in Dulbecco’s modified Eagle medium (DMEM) with high glucose (ThermoFisher) supplemented by 10% Fetal Bovine Serum (FBS; Gibco, ThermoFisher), 1% sodium pyruvate (Gibco, ThermoFisher), 1% L-glutamine (Gibco, ThermoFisher), and 1% Penicillin-Streptomycin (Gibco, ThermoFisher). Cells were cultured in a 10-cm culture dish and incubated at 37^°^C and 5% CO_2_. Cells were passed twice a week at 90% confluence, by washing with PBS, incubating with Trypsin/EDTA (Gibco, ThermoFisher) for 3-5 minutes at 37^°^C, and diluting the cells in fresh medium (1:10) on a new plate.

COS-7 cells were seeded either on 24 mm high-precision cover glasses (Carl Roth) in a 6-well plate or μ-Slide 8 Well high Glass Bottom (Ibidi). The cover glasses were first plasma cleaned by exposure to O_2_-plasma for 2 minutes, making the surface hydrophilic allowing better adhesion of cells for microscopy experiments. Cells at 90% confluence were appropriately diluted at a 1:10 ratio and seeded onto the substrates. Following seeding, the cells were incubated at 37^°^C and 5% CO_2_ overnight followed by fixation procedures.

#### Immunostaining

COS-7 cells were fixed when moderate confluence containing single cells was reached. Generally, this means that the samples were fixed about 24 hours after seeding. Cells were extracted for 90 seconds at room temperature in prewarmed (37^°^C) extraction buffer containing 0.3% (vol/vol) Triton X-100 and 0.25% (wt/vol) Glutaraldehyde in Microtubule-stabilizing buffer Kapitein (MTSBK; 80 mM PIPES, 7 mM MgCl_2_, 1 mM egtazic acid, 150 mM NaCl, 5 mM D-glucose). The extraction buffer was replaced by prewarmed (37^°^C) fixation buffer (4% (wt/vol) paraformaldehyde in PBS) and incubated for 10 minutes at room temperature. The fixation buffer was removed by washing three times with PBS for five minutes under a small traveling wave in each chamber. After fixation, the cell samples were either stored in PBS with 50% (vol/vol) glycerol at 4^°^C for up to three days, or the samples were directly quenched.

Fluorescent quenching was performed by incubating 10 mM freshly prepared sodium borohydride in PBS for 7 minutes at room temperature. This was followed by a quick wash with PBS and two washes of 10 minutes on the orbital shaker. The fixed cells were permeabilized with 0.25% (vol/vol) Triton X-100 in PBS, incubated for 7 minutes at room temperature at the orbital shaker and followed by three washes of 5 minutes on the orbital shaker with PBS. Fixed cells were blocked with blocking buffer (BKK; 2% (wt/vol) bovine serum albumin, 10 mM glycine, 50mM NH_4_Cl) either for 60 minutes at room temperature or overnight at 4^°^C.

Primary and secondary antibodies or nanobodies were diluted in BKK according to the desired degree of labeling. Incubation was done for each of the stainings for 1 hour at room temperature in an incubation chamber and followed by three washes with BKK for 5 minutes on the orbital shaker. After the secondary staining and corresponding washes, the samples were post-fixated by incubating for 10-15 minutes with 2% (wt/vol) paraformaldehyde in PBS. Finally, the fixated cells were washed thrice with PBS for 5 minutes on the orbital shaker. Samples were stored in 50% glycerol in PBS at 4^°^C until used.

### Microscope setup

Microscopic images were captured using a custom-built microscope based on the open microscope frame MiCube^24^. Full details of the microscope setup will be published elsewhere. To provide a brief overview, the setup incorporates a 1 W 638 nm laser (LAB-638-1000, Lasertack), which is combined using a multimode optical fiber (M42L05, Thorlabs). The laser beam is then collimated and directed onto the rear focal plane of an oil-immersion objective (NA 1.5, 60x, UPLAPO60XOHR, Olympus). The system incorporates a one-dimensional motorized stage (KMTS25E/M, Thorlabs) used to translate the excitation beam in the back focal plane in order to adjust the incident angle towards the optical axis. Hence, facilitating the transition between Epi, HILO, and TIRF illumination modalities.

To ensure homogenous illumination across the sample plane, an optical fiber vibration motor (304-111, precision micro-drives) was used to agitate the optical fiber. Fluorescence is decoupled from the excitation beam using a quad-band dichroic mirror (zt405/488/561/640rpc, Chroma) and further filtered by a notch filter (ZET405/488/561/640mv2, Chroma). After passage through a bandpass emission filter (ET706/95m, AHF Analysentechnik AG), the image is focused and captured using an sCMOS camera (BSI Express, Photometrics). Sample positioning is managed via a three-dimensional Stick-Slip piezo stage (CLS5252, Smaract), and image acquisition is performed using μManager 2.0 gamma.

### Image acquisition

Fixed cells were imaged in an imaging buffer containing 500 mM NaCl in PBS with varying imager strand concentrations at room temperature. In experiments exploring the impact of ethylene carbonate (EC), 5% (v/v) EC (Fisher Scientific) was introduced into the imaging buffer.

Microtubule imaging was conducted utilizing TIRF illumination, while HILO illumination was employed for imaging mitochondria. For each experiment, the selection of exposure time was based on a qualitative assessment of blinking kinetics and the SNR per frame in combination with the resulting SOFI results. The specific imaging parameters for each image can be found in supplementary table **??**.

For the large FOV imaging, we used the multi-position acquisition in μManager where we generated 4 by 9 grids, with an overlap of 10% between tiles. 500 frames were recorded for one single tile before moving to the next grid. The grids were stitched together later using the Fiji stitching plugin^25^.

### Data analysis

#### SOFI cross-cumulant analysis

The SOFI calculations were performed using a cross-cumulant-based algorithm available from https://www.github.com/kgrussmayer/sofipackage and implemented in MATLAB R2021b. Constant parameters were chosen allowing for comparison between the imaging buffer conditions. The input image sequence was subdivided into subsequences of 1000 frames each. This subsequent length was chosen to minimize the influence of photobleaching. As a pre-processing step, drift correction based on cross-correlation between the different SOFI subsequences was applied. For post-processing, deconvolution parameters were configured with a PSF approximation of a Gaussian with an FWHM of 4.2 pixels and a total of 10 iterations.

To observe the evolution of SOFI results while decreasing the acquisition time, the SOFI calculations were repeated for each acquisition of the titration series with decreasing number of frames.

#### SMLM

The single-molecule localization microscopy (SMLM) reconstruction was conducted using the ThunderSTORM plugin within FIJI. Default settings were applied for image filtering and the approximate localization of molecule parameters. Sub-pixel localization of molecules was achieved utilizing a PFS Integrated Gaussian approach, with a fitting radius set to 4 pixels and an initial sigma of 1.6 pixels, employing a Weighted Least Squares fitting method.

Visualization of the reconstructed data was facilitated through the use of averaged shifted histograms, magnified at 5.0x with an update frequency of 50 frames. Post-localization, drift correction in the xy plane was performed using cross-correlation methods to ensure accurate spatial alignment. Additionally, single-molecule localizations with uncertainty values exceeding 15 were filtered out to enhance data reliability and precision.

#### Decorrelation analysis

The resolution of the SOFI results was evaluated based on image decorrelation analysis described by Descloux et al.^8^. This algorithm computes spatial resolution within a single image by employing partial phase autocorrelation, which involves the application of a mask filter and the computation of cross-correlation coefficients in Fourier space. To elaborate further, the analysis involves two primary steps. First, a normalized Fourier transform is calculated and subsequently cross-correlated with the original input image in Fourier space using Pearson correlation. Second, this cross-correlation procedure is iteratively executed while the Fourier transform is filtered through a binary circular mask featuring a diminishing radius ranging between 0 and 1.

In this work, the resolution of SOFI results was calculated using the MatLab software from https://github.com/Ades91/ImDecorr. To ensure uniformity throughout the calculations, fixed settings were chosen, taking into account factors such as computational efficiency and precision. The normalized frequencies where the decorrelation curve has to be computed ranges from 0 to 1 with 100 equidistant points within this interval. The number of high-pass filters used to calculate the resolution was set to 20.

### Microtubule cross-sections

An alternative approach to assess the resolution is by evaluating the intensity profile of a cross-section of converging microtubules. A perpendicular line profile was defined across a microtubule, and intensity values were recorded and normalized for each experimental condition (including average intensity profile and higher-order SOFI images). This process necessitated appropriate scaling and considered pixel reduction resulting from SOFI post-processing.

#### Jackknife SNR

SOFI-specific SNR characterization was performed to ensure sufficient image quality using Jackknife resampling^7^. The algorithm was implemented as part of the SOFI cross-cumulant algorithm and is computationally expensive. Consequently, SNR estimation was conducted solely for specific, carefully chosen acquisitions.

Jackknife resampling involves creating *N* new datasets, where *N* corresponds to the number of raw images in the original dataset. Each new dataset is generated by excluding one image from the sequence and is subsequently employed to generate a new SOFI image, resulting in *N* new SOFI images. For every pixel value *I*(*x, y*) in the original SOFI image, *N* new values *I*_*n*_(*x, y*) are produced. These values provide a distribution of possible pixel intensities for that specific pixel location. The variation in these values across the new SOFI images provides insight into the uncertainty associated with the original pixel value. This uncertainty can then be used to calculate the SNR per pixel. The SNR per pixel is defined as:

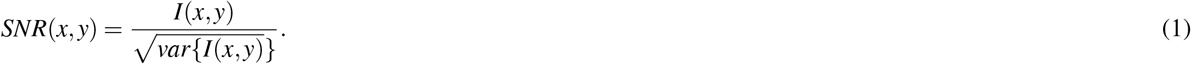

Here, the uncertainty associated with the original pixel value is:

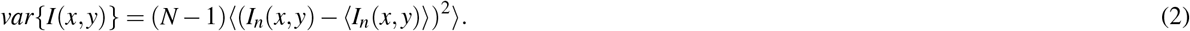

The SNR per pixel is calculated for each subsequence, taking the mean of the SNR values and associated uncertainty yields an SNR value for the final SOFI image with 10,000 frames. Alternatively, to track the change in SNR across the entire raw image sequence, the calculation was performed iteratively by progressively adding frames. The process began with 1,000 frames, and in each subsequent step, an additional 1,000 frames were incorporated.

### Intensity time traces

The methodology employed for intensity analysis involved the computation of normalized average pixel intensities over a temporal sequence. This analysis was conducted within a defined region of interest spanning 3×3 pixels across the entire time series. The selection of the specific pixel area involved the identification of a representative microtubule structure within the average widefield projection image. This strategy aimed to ensure that the chosen region was relevant and reflective of the underlying sample characteristics.

## Acknowledgements

We thank N. van Vliet for cell culture work and R. van der Valk for guidance in protein production. We thank Y. Deurloo for initial protein preparations. We are grateful to all members of the Grussmayer and Geertsema lab for discussions.

This work was supported by the TU Delft Department of Bionanoscience and the Department of Imaging Physics, the TU Delft Bioengineering Institute through an MSc Project Proposal, and the TU Delft AI initiative through BIOLab.

## Author contributions

K.S.G. and H.J.G. conceived the idea, initiated and supervised the project. K.S.G. designed the experiments and helped with data analysis. H.L.W. labeled initial proteins, prepared and imaged initial cell samples and analyzed them. M.N.F.H. and B.K.Z. produced the proteins and performed the conjugations. R.H., and B.K.Z. prepared and imaged cell samples. R.H., M.T. and B.K.Z. performed the data analysis. R.H. built and maintained the microscope used for data collection. K.S.G., H.J.G., R.H., and B.K.Z. wrote the manuscript with input from all authors. All authors discussed the results and commented on the manuscript..

