## Supplementary Information for "High-throughput single molecule microscopy with adaptable spatial resolution using exchangeable oligonucleotide labels"

### ABSTRACT

### Supplementary information

#### Supplementary note 1: SOFI theory

Super-resolution optical fluctuation imaging (SOFI) is a super-resolution technique that enhances spatial resolution by computing higher-order statistics of a time series of stochastically blinking fluorophores. SOFI does not demand major technical modifications to the microscope, e.g. a standard widefield microsocpe can be used, and is flexible regarding fluorophores and imaging conditions. The only requirement is that the sample has been labeled with fluorescent emitters that switch between two (or more) distinct states independently from each other and repeatedly in a stochastic way[1].

SOFI's super-resolving power stems from the assumption that fluorescent emitters switch independently and stochastically, correlating only in time and space with themselves. This independence, combined with the mathematical properties of cumulants (statistical quantities related to correlation functions), enables the extraction of higher-order statistical information from the blinking patterns of the fluorophores. The cumulants effectively filter out cross-terms and uncorrelated noise, allowing for the reconstruction of images with enhanced spatial resolution.

##### *Fluorescent signal and cumulants*

The detected intensity on the camera, assuming $N$ single, independently fluctuating emitters, can be described by:

$$I(\mathbf{r},t) = \sum_{k=1}^{N} U(\mathbf{r}-\mathbf{r}_k)\cdot \varepsilon_k \cdot s_k(t) + b(\mathbf{r},t). \tag{1}$$

where $U(\mathbf{r})$ represents the optical systems point spread function (PSF), and $b(\mathbf{r},t)$ describes the background signal, including uncorrelated noise. The time-dependent molecular brightness is described for each emitter by $\varepsilon_k \cdot s_k(t)$, where $\varepsilon_k$ denotes the constant molecular brightness and $s_k(t)$ represents a time-dependent fluctuation component with values of 0 (off-state) and 1 (on-state).

To capture detailed patterns of fluorescence signal fluctuations which contribute to the super-resolved image, we analyze nth-order cumulants. In general, spatio-temporal cumulants can be calculated using various time lags and different combinations of pixels to explore both temporal and spatial correlations in the fluorescence signal. We use a time lag of zero to simplify the analysis and cross-cumulants between different pixels to suppress shot noise[2–4]. Our SOFI algorithm generates virtual pixels which are essentially interpolated points between the physical pixels captured by the camera. The higher-order cross-cumulants thus inherently provide a finer sampling of the super-resolved image. Mathematically, the $n^{th}$-order cumulant with zero time lag can be written as:

$$\kappa_n\{I(\mathbf{r},t)\} = \sum_{k=1}^{N} U^n(\mathbf{r}-\mathbf{r}_k)\varepsilon_k^n \kappa_n\{s_k(t)\} + \kappa_n\{b(\mathbf{r},t)\}. \tag{2}$$

In the context of SOFI, the fluorophore's blinking behavior, $s_k(t)$, can be thought of as a binary function taking the value 1 when the fluorophore emits and 0 when it is off. Its cumulant, $\kappa_n\{s_k(t)\}$, is a Bernoulli distribution with probability $\rho$ (i.e. on-time ratio) and can be approximated by an $n^{th}$-order polynominal function of the on-time ratio $f_n(\rho) = (1-\rho)\frac{\partial f_n}{\partial \rho}$. Moreover, the term $\kappa_n\{b(\mathbf{r},t)\}$ for a constant background and additive noise is eliminated from higher-order cumulants because additive noise is assumed random and uncorrelated. Hence, we can estimate the $n^{th}$-order cumulant with zero time lag as:

$$\kappa_n\{I(\mathbf{r},t)\} = \varepsilon_k \cdot f_n(\rho) \sum_{k=1}^{N} U^n(\mathbf{r}-\mathbf{r}_k) \cdot \varepsilon_k^n. \tag{3}$$

##### *Resolution enhancement*

The effective PSF in SOFI is raised to the power of $n$, leading to a narrowing of the PSF by a factor of $\sqrt{n}$, consequently resulting in enhanced spatial resolution. Using deconvolution techniques enables a final resolution improvement that scales linearly with the order of the cumulant. However, higher-order SOFI imaging comes with challenges, including the amplification of heterogeneities in molecular brightness $\varepsilon^n$, which may mask less bright structural details. To address this issue, Geissbuehler et al. proposed deconvolving the cumulant images and then linearizing the brightness response by applying the n-th root to the deconvolved cumulant image[2,5].

It is important to note that while there are no theoretical limitations to calculating higher SOFI, the signal-to-noise ratio and signal-to-background ratio typically decrease, increasing the risk of processing artifacts. Additionally, computational complexity grows with the order $n$, leading to higher memory requirements and longer processing times.

#### Supplementary note 2: Buffer optimization

To accelerate both binding and unbinding rates in our experiments, we optimized the imaging buffer composition by increasing the imager strand concentration and incorporating ethylene carbonate (EC). This Supplementary Note details the buffer optimization process and its impact on the blinking kinetics.

. Our approach is supported by On-time refers to when the imager strand is bound, while off-time refers to when it is unbound. This indicates that optimizing the blinking kinetics is critical for enhancing SOFI imaging quality

Microtubules in fixed COS-7 cells were immunostained with a speed optimized docking strand[6]. By increasing the imager strand concentration, the probability of hybridization of the DNA oligos is raised (Fig 1a,b). Compared to the initial imager strand concentration (Fig 1a), an increase in signal per frame is observed, indicating more frequent binding upon addition of imager strands. This is confirmed by evaluating the intensity time traces over time (Supplementary Fig 2).

As for any fluorescence imaging, the SOFI signal contrast depends on the SNR and signal to background ratio (SBR), but also on the fluctuation statistics and sampling of the blinking. Solely increasing the imager strand concentration has limitations for high-order SOFI reconstructions, as it raises background noise, reducing the SNR and SBR. Moreover, when the off-time is reduced to the point of continuous occupation of resolvable binding sites, fewer intensity fluctuations are observed, consequently diminishing the SOFI signal. Cevoli et al. performed a parameter optimization for SOFI using a design of experiments approach, where simulated data was used to investigate the parameters influencing the quality of SOFI images[7]. The analysis indicated that the best SOFI results were achieved when the fluorescent on- and off-times were minimized. Thus, to obtain more intensity fluctuations the on-time should also be minimized.

Prior research has demonstrated the acceleration of DNA dehybridization through the incorporation of ethylene carbonate (EC)[8]. This aprotic solvent increases the solubility of the DNA bases and has been effectively employed to accelerate DNA-PAINT experiments by decreasing the on-time without affecting the off-time[9]. Supplementing the high imager strand concentration buffer with EC leads again to the appearance of single blinks, signifying a notable reduction in on-time (Fig 1c ). This observation is confirmed by comparing the intensity time traces over time: upon the addition of EC, single peaks are observed whereas without EC, the intensity fluctuations in the signal are less apparent (Supplementary Fig 2). Further increasing the concentration of EC is eventually hindered due to changes in the refractive index of the buffer (data not shown).

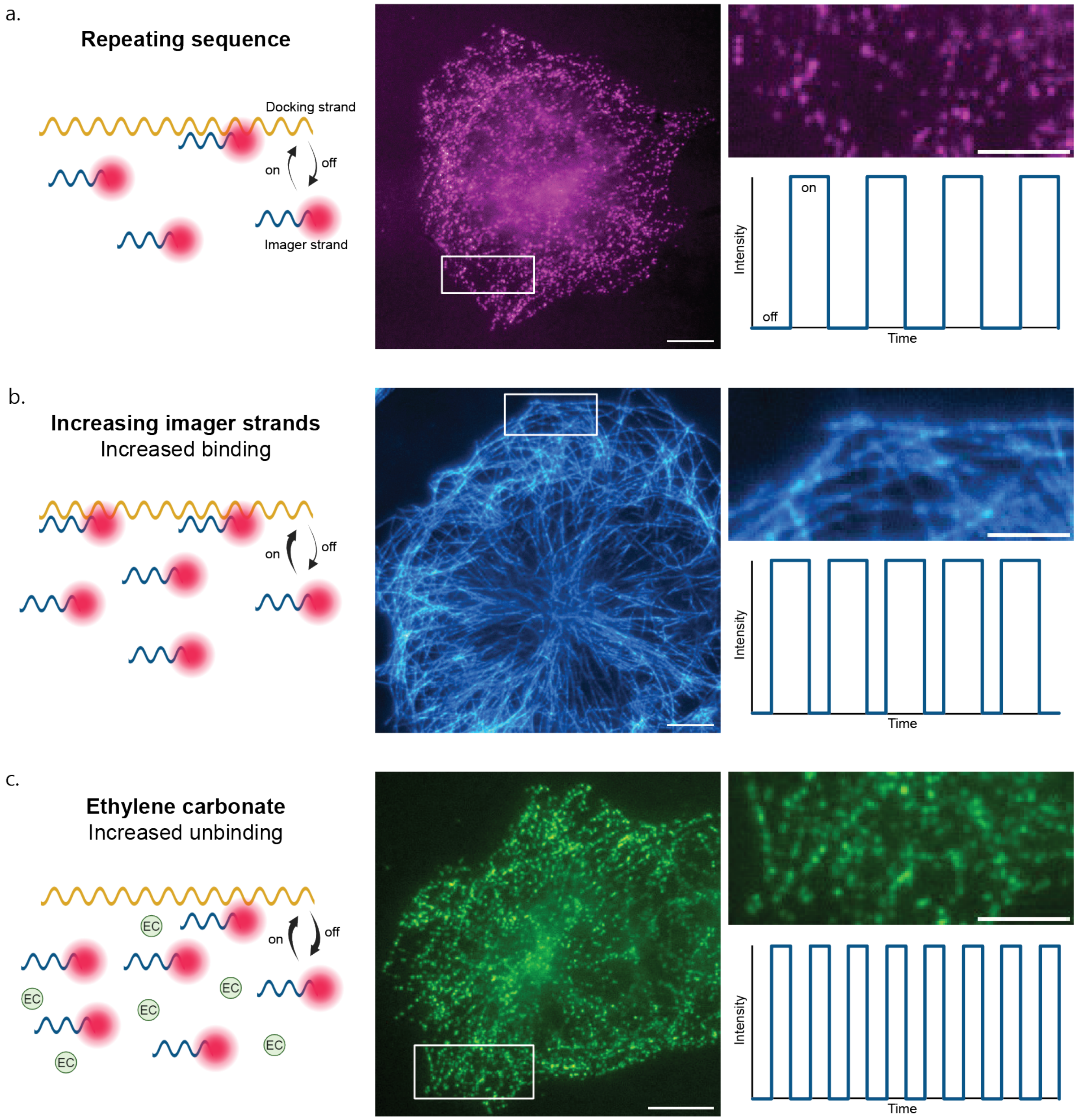


**Figure 1. Influence of buffer composition on blinking kinetics.** (a) Raw image frame displaying blinking behavior under low concentration of imager strands of COS7 cells stained for microtubules with repeating docking sequence[6] (scale bar 10 $\mu$m). Close up of indicated area (scale bar 5 $\mu$m) and representation of corresponding blinking intensity time trace. (b) Blinking behavior with increased imager strand concentration yielding an increased binding frequency. (c) Blinking behavior of increased imager strand concentration and addition of ethylene carbonate yielding both an increased binding and unbinding frequency.

| Figure | Docking strand | Buffer composition: [Imager strand], [EC] | Frames | Exposure time (ms) | Illumination intensity (kW $cm^{-2}$) |
|---|---|---|---|---|---|
| Figure?? | 5xR1 | 100 pM | 10,000 | 30 | 2.5 |
| Figure ??a & Supplementary figure5a | 5xR1 | 20 nM, 5% EC | 500 | 10 | 2.5 |
| Figure ??b & Supplementary figure5b | 5xR1 | 20 nM, 5% EC | 5,000 | 10 | 2.5 |
| Figure ??c & Supplementary figure5c | 5xR1 | 50 pM | 30,000 | 100 | 3.1 |
| Figure ??a | 5xR1 | 20 nM, 5% EC | 500 | 10 | 1.2 |
| Figure ?? SOFI 2 | 5xR1 | 5 nM, 5% EC | 2,500 | 30 | 3.1 |
| Figure ?? SOFI 6 | 5xR1 | 5 nM, 5% EC | 8,000 | 30 | 3.1 |
| Figure ?? SMLM | 5xR1 | 25 pM | 15,000 | 100 | 3.1 |
| Figure 1a | 5xR1 | 100 pM | 10,000 | 50 | 2.5 |
| Figure 1b | 5xR1 | 5 nM | 10,000 | 10 | 2.5 |
| Figure 1c | 5xR1 | 5 nM, 5% EC | 10,000 | 30 | 3.1 |
| Supplementary figure 3 | 5xR1 | 5 nM, 5% EC | 10,000 | 50 | 3.1 |

**Table 1.** Summary of samples, buffer, and imaging parameters

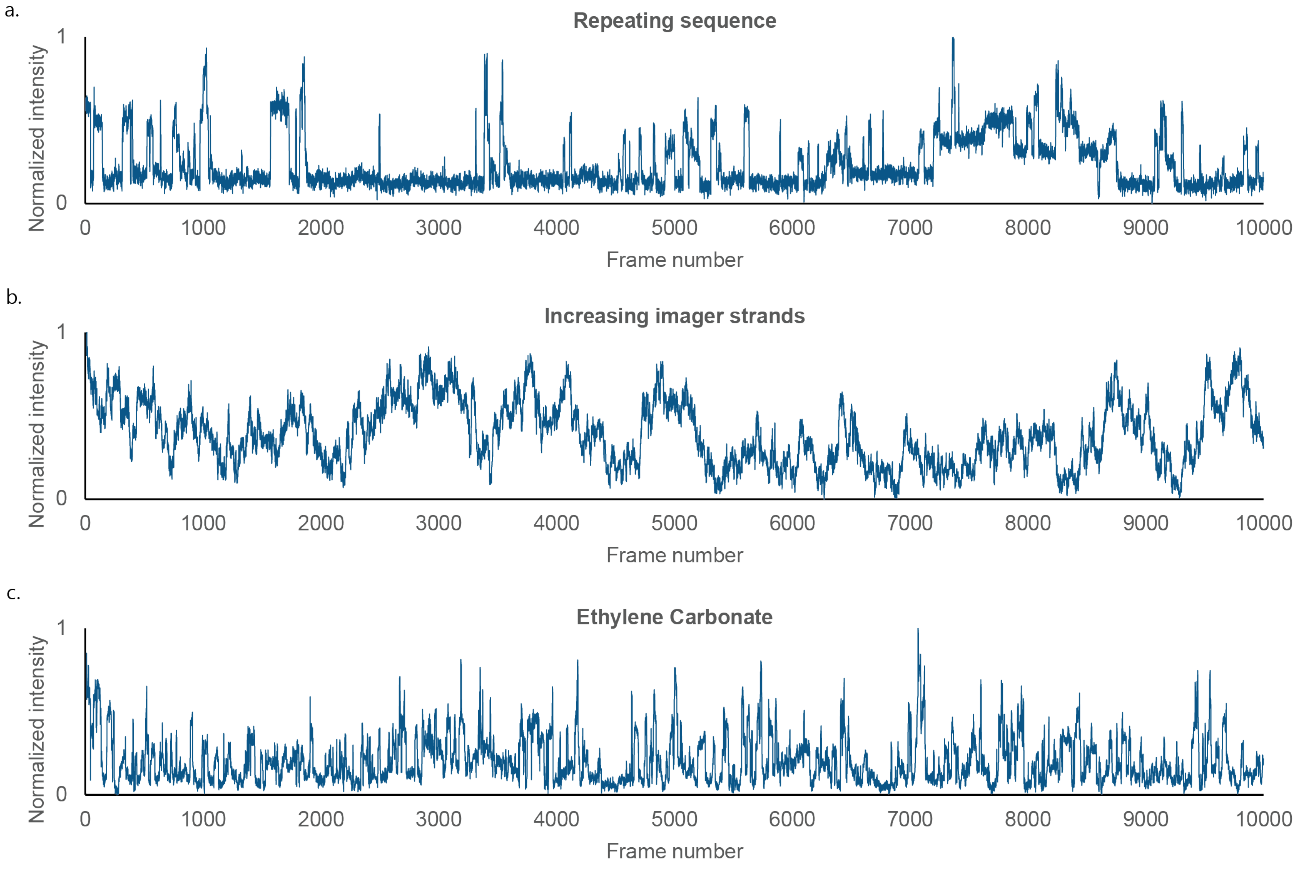


**Figure 2. Intensity time traces** (a) 100 pM (b) 5 nM (c) 5nM + EC.

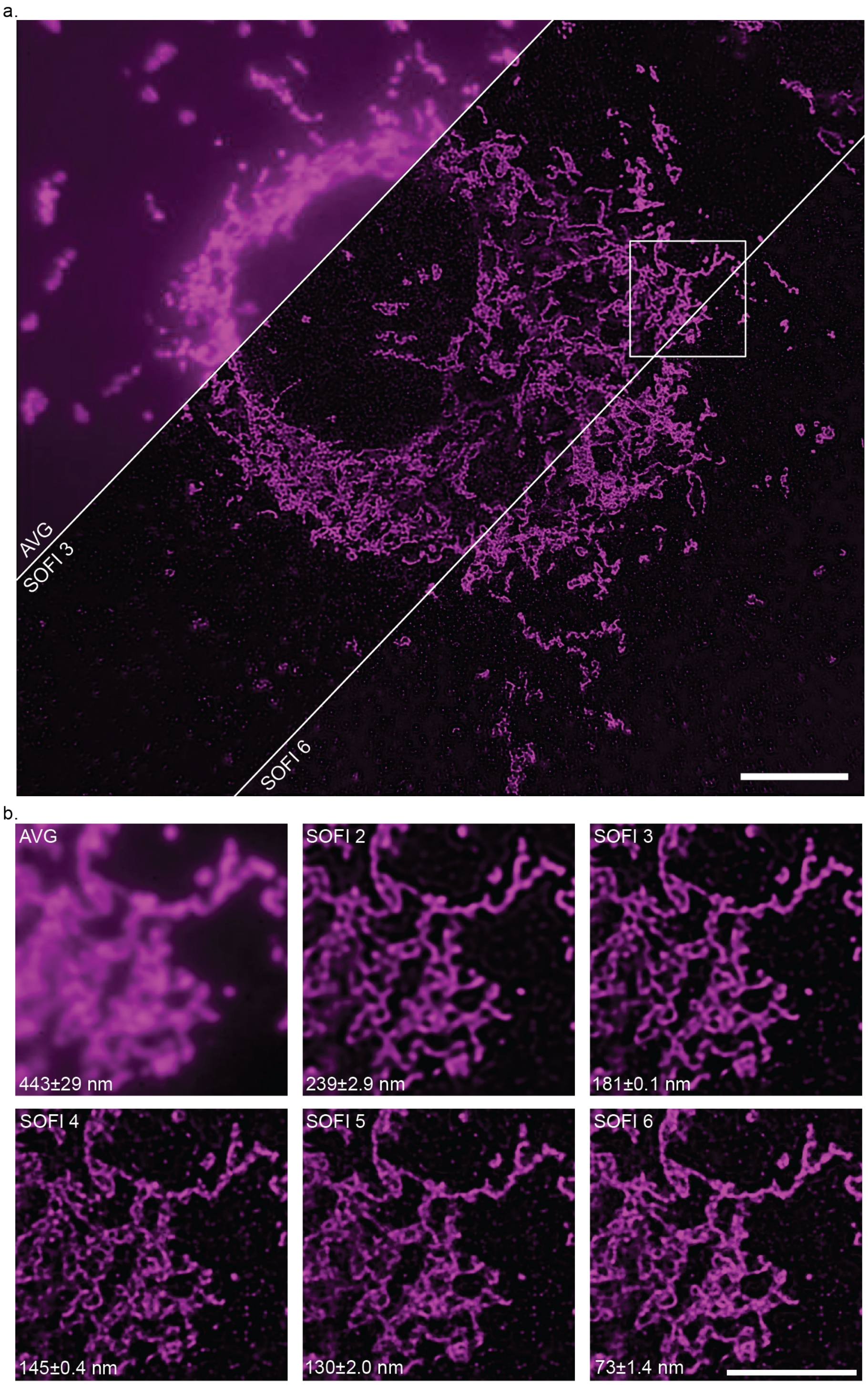


**Figure 3. SOFI reconstructions up till 6th order** (a) sc 10um (b) zoom ins, sc 5um

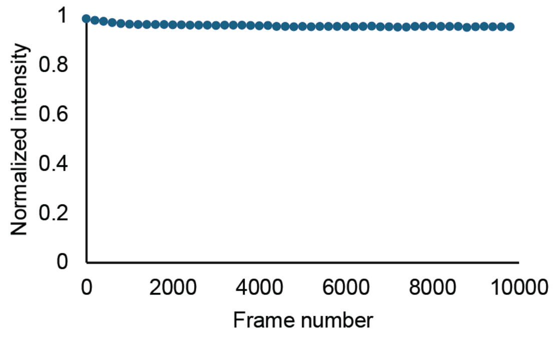


**Figure 4.** **Mean fluorescence signal over time** Mean fluorescence signal of full time series of 3 COS7 cells stained for microtubules with DNA-PAINT.

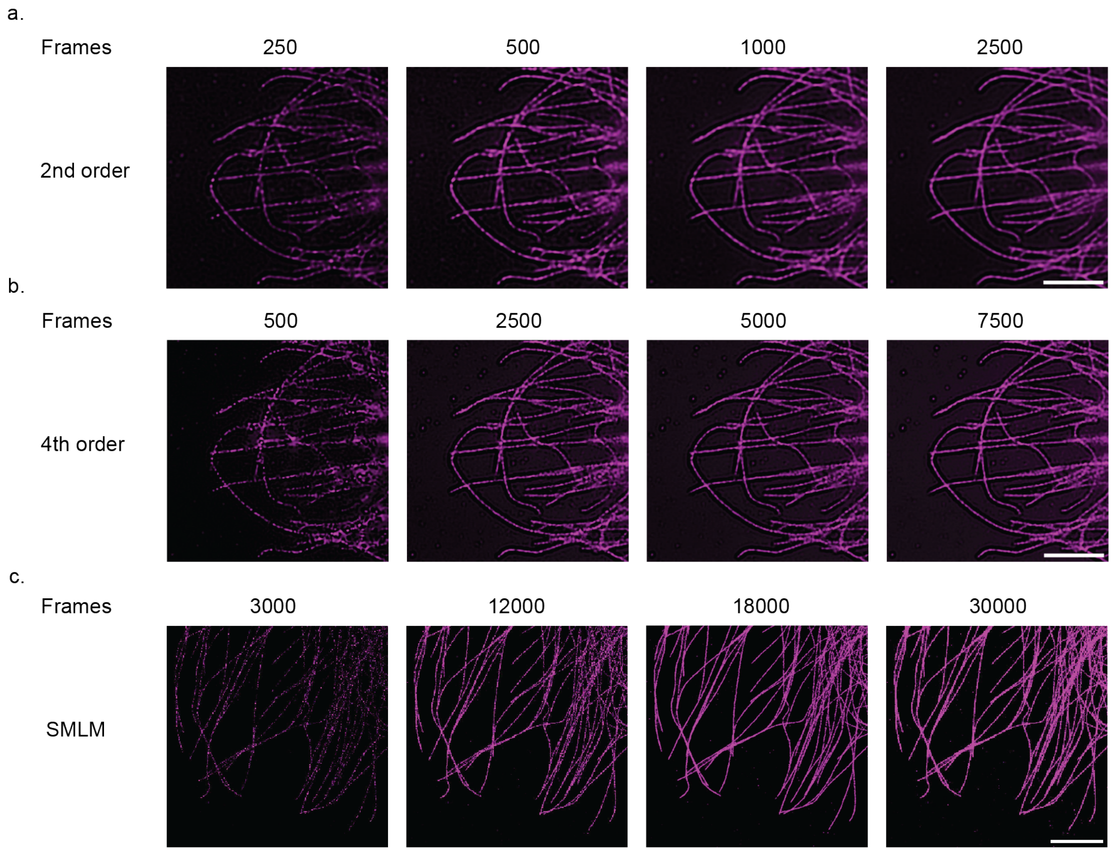


**Figure 5.** **Evaluation of SOFI and SMLM signal with extending acquisition time**
